# Phosphorylation of Aly3 C-terminus impedes aberrant endocytosis of *S. pombe* hexose transporter Ght5

**DOI:** 10.1101/2024.03.24.586504

**Authors:** Yusuke Toyoda, Fumie Masuda, Shigeaki Saitoh

## Abstract

In fission yeast, *Schizosaccharomyces pombe*, transcriptional upregulation and cell-surface localization of the hexose transporter, Ght5, are required for cell proliferation in low glucose. As the target of rapamycin complex 2 (TORC2) signaling pathway inhibits α-arrestin Aly3-dependent endocytosis of Ght5, we hypothesized that this endocytosis was inhibited by phosphorylation. To identify phosphorylation sites required for cell proliferation in low glucose, serine and threonine residues of Aly3 and Ght5 reportedly phosphorylated were replaced with alanine. We found that C-terminal serine residues of Aly3, but not Ght5, are necessary for proliferation in low glucose. Expression of Aly3 protein unphosphorylated at the C-terminus led to increased ubiquitination and vacuolar accumulation of Ght5 in low glucose, but reversion of one of the alanine residues to serine reduced ubiquitination and vacuolar accumulation of Ght5. Also, Aly3 physically interacted with the HECT-type ubiquitin ligases Pub1 and Pub3, and these interactions were required for surface localization of Ght5 and proliferation in low glucose. This study reveals mechanisms by which Aly3 is regulated so that fission yeast can adapt to nutritional stress.

## Introduction

Eukaryotic cells need to acquire nutrients via transporters and channels expressed on the cell surface so that they can grow and proliferate. While cells are able to adapt to changes in the nutritional environment, molecular mechanisms underlying such adaptation are only partially understood. Whereas *Schizosaccharomyces pombe* cells are usually cultivated in the laboratory in medium containing high concentrations (2–3%) of glucose, they are able to proliferate readily in medium with concentrations as low as 0.08% (Saitoh and Yanagida, 2014). Among the 8 hexose transporter genes (*ght1*^+^ to *ght8*^+^), only *ght5*^+^, which encodes a high-affinity hexose transporter is essential for *S. pombe* cells to proliferate in low-glucose, whereas it is not essential in high-glucose (Heiland et al., 2000; Wood et al., 2002; Saitoh et al., 2015). To adapt to glucose limitation, fission yeast cells regulate both the expression level and subcellular localization of Ght5 (Toyoda and Saitoh, 2015). Upon glucose limitation, transcription of *ght5*^+^ is upregulated by mobilizing the transcription repressor Scr1 from the nucleus to the cytosol (Saitoh et al., 2015). This process is dependent on Ca^2+^/calmodulin-dependent kinase kinase (CaMKK) and AMP-dependent protein kinase (AMPK). In abundant glucose, Ght5 is expressed at a low level and localizes to the cell surface predominantly near the cell equator, whereas in low glucose, Ght5 is expressed over the entire cell surface (Saitoh et al., 2015). Intriguingly, limitation of nitrogen sources, e.g. NH_4_Cl and amino acids, but not glucose, promotes transport of Ght5 from the cell surface to vacuoles (organelles equivalent to lysosomes in higher eukaryotes), regardless of the glucose concentration in the medium (Toyoda et al., 2021), suggesting overlapping control of glucose uptake by nitrogen availability. Controlling localization of Ght5 during nutritional stress is critical, as its failure leads to defective cell proliferation and/or decreased viability.

Target of rapamycin complexes 1 and 2 (TORC1 and TORC2) govern cellular responses to nutritional stress. TORC1 and TORC2 are evolutionarily conserved, responding to changes in nutrient availability and modulating cellular functions by phosphorylating downstream AGC-family protein kinases (Wullschleger et al., 2006). In *S. pombe*, TORC1 is composed of a catalytic Tor2 subunit and accessory subunits, including Mip1/Raptor, whereas TORC2 is composed of a catalytic Tor1 subunit and accessory subunits, including Ste20/Rictor (Hayashi et al., 2007; Tatebe and Shiozaki, 2017). TORC1 senses growth factors, energy levels, and nitrogen sources, and promotes cell growth and biosynthesis of proteins, lipids and nucleotides, but it inhibits autophagy (Otsubo and Yamamato, 2008; Stracka et al., 2014; Saxton and Sabatini, 2017). On the other hand, in fission yeast, TORC2 senses availability of glucose and nitrogen sources, and regulates stress responses, sexual development, metabolism, and cell proliferation (Kawai et al., 2001; Weisman et al., 2001; Jacinto and Lorberg, 2008; Hatano et al., 2015; Toyoda et al., 2021). Unlike TORC1, TORC2, its downstream Akt-related kinase Gad8 and the phosphoinositide-dependent protein kinase 1 (PDK1)-like kinase Ksg1, which comprise the TORC2 signaling pathway (Cybulski and Hall, 2009), are required for cell-surface localization of Ght5 under low-glucose (Saitoh et al., 2015). Mutants deficient in the TORC2 pathway exhibit vacuolar localization of Ght5 and defective proliferation in low glucose, whereas these defects are not observed in mutants defective in TORC1. These defects of the *gad8*^ts^ mutant were suppressed by mutations of genes encoding one of the α-arrestins, Aly3, and subunits of endosomal sorting complexes required for transport (ESCRTs) (Toyoda et al., 2021). α-arrestins are thought to promote ubiquitination of specific membrane proteins by recruiting the HECT (homologous to the E6-AP carboxyl terminus)-type ubiquitin ligase (Puca and Brou, 2014). Once endocytosed, ubiquitinated cargos are recognized by the ESCRT-0 complex, and the ESCRT-I, II and III complexes successively process and enclose them in endosomes, forming multivesicular bodies (MVBs), which are transported to vacuoles and degraded (Schmidt and Teis, 2012). Ubiquitination of Ght5 is increased in TORC2 pathway mutants in low glucose in an Aly3-dependent manner (Toyoda et al., 2021). Therefore, our previous results indicate that the TORC2 pathway inhibits Aly3-dependent ubiquitination and vacuolar transport of Ght5 in low-glucose. However, it is not clear how Aly3 activity is controlled.

Arrestins, originally identified as proteins that attenuate signaling mediated by rhodopsin, a G-protein-coupled receptor (GPCR) expressed in retinal cells (Wilden et al., 1986; Zuckerman and Cheasty, 1986), comprise a large protein family, homologs of which are found in archaea, bacteria, and eukaryotes (Alvarez, 2008). In eukaryotes, three types of arrestins, α-arrestins, visual/β-arrestins, and VPS26-like proteins, are known (Alvarez, 2008). The human genome encodes 6 α-arrestins, 4 β-arrestins, and 4 VPS26-like proteins, whereas budding yeast, *Saccharomyces cerevisiae,* expresses 14 α-arrestins and one VPS26 protein (Alvarez, 2008; Kahlhofer et al., 2021). According to PomBase, the fission yeast database (Harris et al., 2022), the *S. pombe* genome encodes 11 arrestin-like proteins, including 4 α-arrestins. α-arrestins typically have Arrestin-N and Arrestin-C domains (InterPro ID: IPR011021 and IPR011022, respectively (Paysan-Lafosse et al., 2023)) at the N-terminus and multiple proline-rich PY motifs at the C-terminus (Alvarez, 2008). It is thought that the arrestin domain interacts with membrane proteins, and that the PY motif interacts with the WW domain of HECT-type ubiquitin ligases (Andoh et al., 2002; Kahlhofer et al., 2021). Through these interactions, α-arrestins behave as ubiquitin ligase adaptors for a specific set of transporter proteins, by which cells regulate their cell surface proteomes to adapt to nutritional stress. Selectivity of α-arrestins for transporter proteins appears high. Among four α-arrestins in fission yeast (Aly1, Aly2, Aly3 and Rod1), only Aly3 regulates surface localization of Ght5 (Toyoda et al., 2021). Also, surface localization of budding yeast hexose transporters is regulated only by certain α-arrestins, including Rod1/Art4 and Rog3/Art7, which are homologous to fission yeast Aly3 (Kahlhofer et al., 2021).

Post-translational modification is a major means of regulating arrestins. Phosphorylation is thought to inactivate α-arrestins by promoting interaction with 14-3-3 proteins in budding yeast or by enhancing proteasome-dependent degradation in mammals (Zhang et al., 2010; Llopis-Torregrosa et al., 2016; O’Donnell and Schmidt, 2019). TXNIP, a mammalian homolog of Aly3, is inactivated and retracted from the cell surface when phosphorylated by AMPK or Akt at serine 308 (S308) (Wu et al., 2013; Waldhart et al., 2017). Rod1 is phosphorylated by AMPK and inactivated by binding to 14-3-3 proteins (Shinoda and Kikuchi, 2007; Llopis-Torregrosa et al., 2016). Conversely, dephosphorylation is thought to activate α-arrestins and to promote selective endocytosis of transporter proteins. In budding yeast, phosphatases, including Glc7 (PP1), Sit4 (PP2A) and calcineurin (PP2B), reportedly dephosphorylate α-arrestins (Kahlhofer et al., 2021). Ubiquitination at K263 of the fission yeast arrestin-related protein Arn1/Any1 is necessary for interaction with the HECT-type ubiquitin ligase, Pub1 (Nakashima et al., 2014). Ubiquitination of budding yeast Cvs7/Art1 by the HECT-type ubiquitin ligase Rsp5 is required for endocytosis of the arginine transporter, Can1 (Lin et al., 2008). However, mechanisms that regulate functions of α-arrestins are not fully understood.

In this study, we sought to discover the molecular mechanism by which Aly3 is negatively regulated under low glucose. Cells expressing phosphorylation-defective Aly3 failed to proliferate in low glucose. Phosphorylation sites responsible for cell proliferation in low glucose are located at the C-terminus, and phosphorylation of Aly3 at the C-terminus represses ubiquitination of Ght5, consequently ensuring cell proliferation in low glucose by maintaining cell-surface localization of Ght5. Kinase(s) phosphorylating the Aly3 C-terminus and molecular mechanisms regulating physical interaction between Aly3 and Ght5 by phosphorylation are discussed.

## Results

### Fission yeast cells expressing phosphorylation-defective Ght5 proliferate in low glucose

Previous reports suggest that the TORC2 pathway inhibits Aly3 to ensure cell-surface localization of Ght5 and cell proliferation in low glucose (Saitoh et al., 2015; Toyoda et al., 2021). As the TORC2 pathway consists of a cascade of kinases, we hypothesized that subcellular localization of Ght5 is controlled via phosphorylation. Loss of such phosphorylation would impact cell proliferation in low glucose, similarly to the TORC2 pathway-defective mutants, *gad8*^ts^ and *tor1*Δ. Target protein(s) phosphorylated in response to glucose shortage could include Ght5 or proteins required for endocytosis of Ght5, such as Aly3. First, we tested whether phosphorylation of Ght5 is required for cell proliferation in low glucose. Ght5 is reportedly phosphorylated at 11 residues, all of which are located in the C-terminal cytoplasmic tail of approximately 100 residues (Kettenbach et al., 2015). To examine the possibility that phosphorylation of these residues is required for surface localization of Ght5, we constructed a mutant allele of the *ght5*^+^ gene, *ght5*(ST11A), in which all 11 of these serine /threonine residues were mutated to alanine, and inserted this allele into the genome of a *ght5* gene-deleted strain (*ght5*Δ). For control strains, the empty vector or that containing the wild-type allele of the *ght5* gene (*ght5*^+^), was inserted. In the resulting strains, the inserted *ght5* gene was located proximal to the *nmt1*^+^ gene and transcribed from the *nmt1* promoter, which is repressed by thiamine and induced by its removal (Maundrell, 1990). These cells were spotted on high- and low-glucose solid EMM2 minimal media with and without thiamine, and their proliferation was observed (**Fig. 1** and **Supplementary Fig. 1A**). All strains proliferated in high glucose (2%) with or without thiamine. Vector-integrated control cells, equivalent to *ght5*Δ, failed to proliferate on low-glucose media (0.2 and 0.14%), regardless of the presence of thiamine. In contrast, cells expressing the *ght5*^+^ and *ght5*(ST11A) genes proliferated to form colonies on low-glucose media lacking thiamine. Sizes and densities of colonies were identical in both *ght5*^+^ and *ght5*(ST11A) cells. This result indicates that phosphorylation of Ght5 is dispensable for cell proliferation in low glucose.

**Figure 1.**
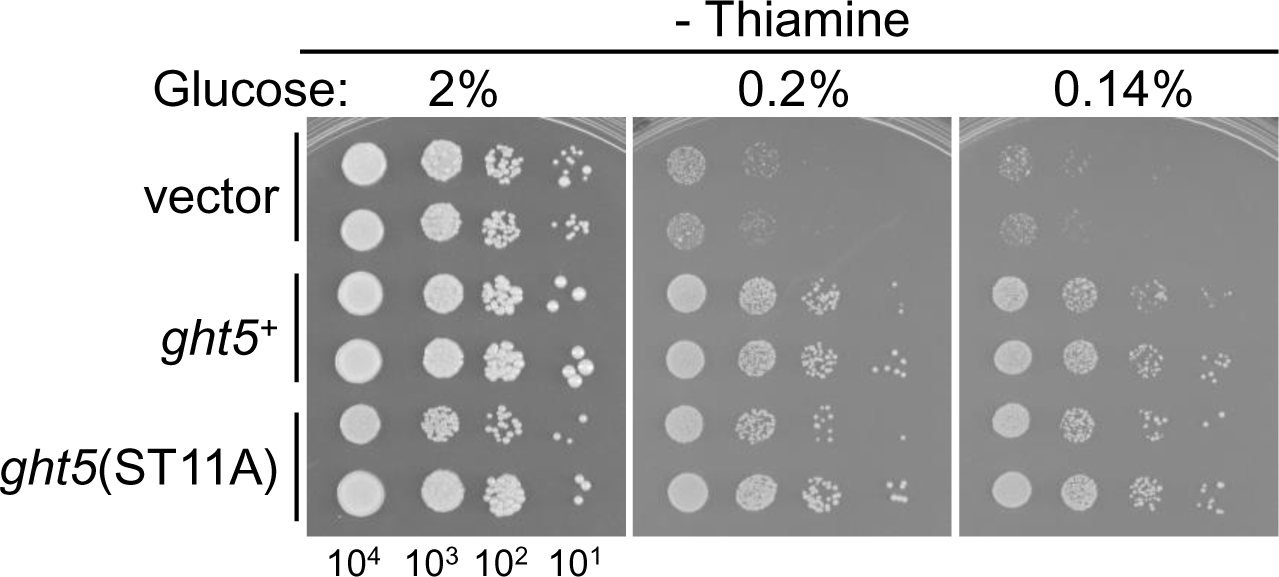
Cells expressing phosphorylation-defective Ght5 protein proliferate in low glucose. Aliquots of *ght5*Δ cells expressing the vector or the wild-type and phosphorylation-defective *ght5* genes were serially diluted 10-fold, spotted onto solid EMM2 medium containing the indicated concentrations of glucose, but not thiamine, and cultivated for 5 days at 33°C. For each strain, two independent clones were tested. At the bottom, the numbers of cells spotted are shown. See **Supplementary Figure 1A** for proliferation of cells in the presence of thiamine.

### Phosphorylation-defective mutation in *aly3* abolishes cell proliferation in low glucose

We then hypothesized that phosphorylation of Aly3 might be required for cell proliferation in low glucose. Fission yeast Aly3 is reportedly phosphorylated at 18 serine and threonine residues (Kettenbach et al., 2015; Tay et al., 2019; Halova et al., 2021). To test this hypothesis, we produced a plasmid harboring a mutant *aly3* gene, *aly3*(ST18A), in which all 18 serine and threonine residues subject to phosphorylation were replaced with alanine (**Fig. 2A**), and made a *S. pombe* strain expressing the mutant *aly3* gene from the *nmt1* promoter by inserting the linearized plasmid into the genome of the *aly3Δ* mutant by homologous recombination (see **Materials and Methods** for details). As a control, a *S. pombe* strain expressing the wild-type *aly3^+^*gene from the *nmt1* promoter was generated in a similar manner. Fixed numbers of cells were spotted onto solid EMM2 media containing different concentrations of glucose with and without thiamine (**Fig. 2B**). All strains proliferated comparably on high-glucose (2%) medium lacking thiamine. However, cells expressing *aly3*(ST18A) did not proliferate at all in low glucose (**Fig. 2B**), whereas cells expressing the wild-type *aly3*^+^ gene proliferated. This indicates that at least some of the 18 residues of Aly3, which are reportedly subject to phosphorylation, are essential for Aly3 to ensure cell proliferation under low glucose.

**Figure 2.**
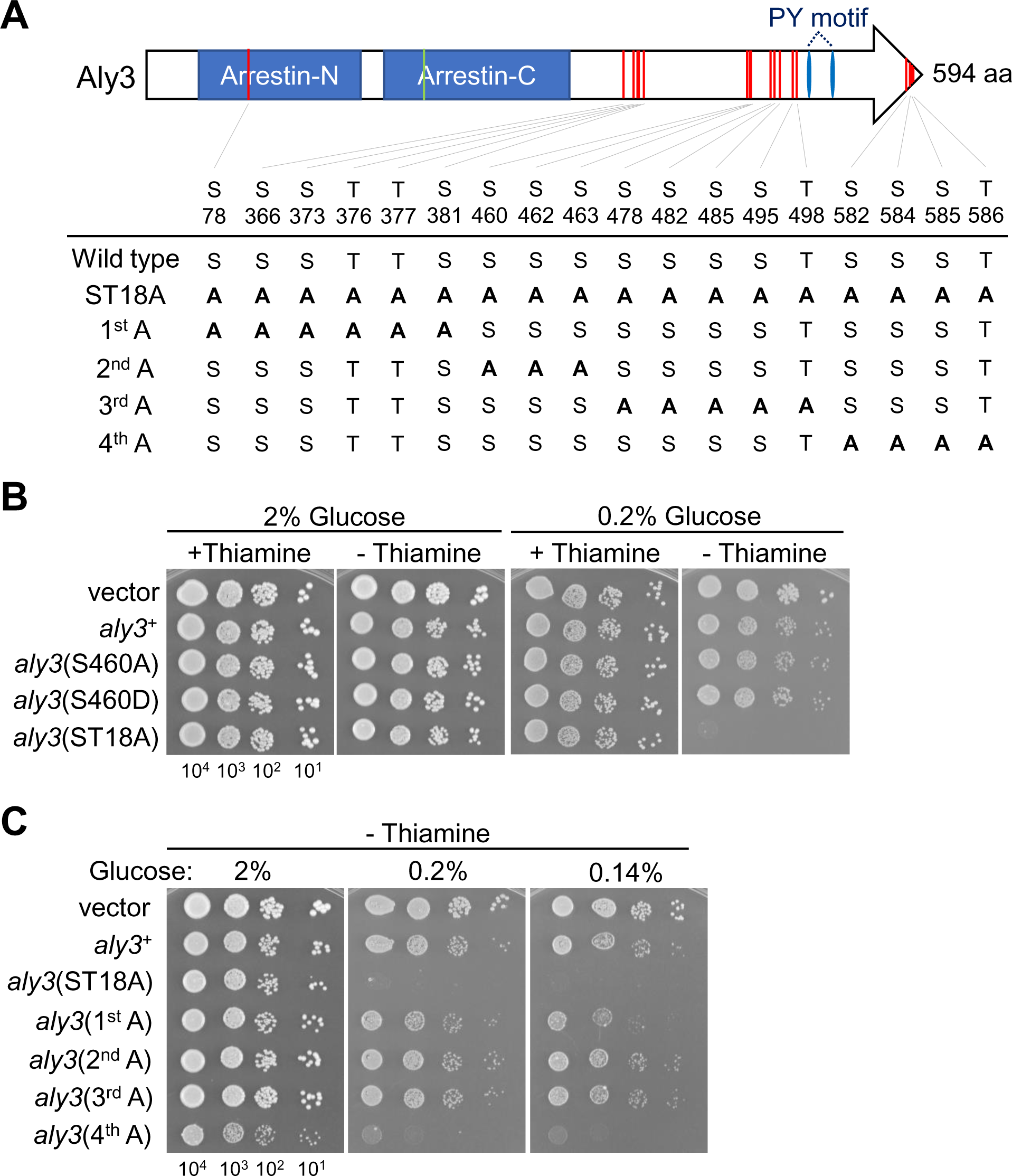
C-terminally located serine and threonine residues of Aly3 are required for cell proliferation in low glucose. **A.** Schematic drawing of domains and post-translationally modified residues of Aly3. Arrestin-N and Arrestin-C indicate the ‘Arrestin-like, N-terminal domain’ and ‘Arrestin C-terminal-like domain’, respectively. Blue vertically elongated ellipse, proline-rich PY motif. Red and green vertical lines indicate phosphorylated and ubiquitinated (K215) residues, respectively (Beckley et al., 2015; Kettenbach et al., 2015; Tay et al., 2019; Halova et al., 2021). At the bottom, amino acid sequences of the wild-type and the phosphorylation-defective Aly3 proteins are shown. **B-C.** Aliquots of *aly3*Δ cells expressing the vector or the wild-type and indicated mutant *aly3* genes were diluted 10-fold, spotted onto EMM2 solid media containing a high (2%) or low (0.2%) concentration of glucose, but not thiamine and cultivated for 5 days at 33°C. At the bottom, numbers of cells spotted are shown. See **Supplementary Figure 1B** for proliferation of cells shown in panel **C** in the presence of thiamine.

Among the 18 reported phosphorylation sites, serine 460 (S460) has a flanking amino acid sequence (R-T-R-A-N-S_460_) that is homologous to the consensus sequence for substrates of Akt kinase (R-x-R-x-x-S/T, where ‘x’ is any amino acid) (Obata et al., 2000). Since *S. pombe* Gad8, which functions downstream of the TORC2 pathway is orthologous to Akt, we predicted that phosphorylation of this serine residue by Gad8 would be essential for cell proliferation in low glucose. Contrary to our prediction, cells expressing Aly3(S460A) or Aly3(S460D), in which S460 was replaced with alanine or aspartate, respectively, proliferated in low glucose comparably to those expressing wild-type Aly3 (**Fig. 2B**), indicating that phosphorylation of Aly3 at S460 is dispensable for cell proliferation in low glucose.

### A cluster of serine residues at the C-terminus of Aly3 is required for cell proliferation in low glucose

The above results suggest that these 18 residues include phosphorylation sites of Aly3 required for cell proliferation in low glucose. To identify such phosphorylation sites, the 18 Ser/Thr residues were divided sequentially into 4 groups from the N-terminus, and all Ser/Thr residues in a group were replaced with alanine for construction of four mutant alleles of the *aly3* gene: *aly3*(1^st^ A), *aly3*(2^nd^ A), *aly3*(3^rd^ A) and *aly3*(4^th^ A) (**Fig. 2A**). Fission yeast cells expressing these mutant *aly3* genes driven by the *nmt1* promoter from the genome were produced by homologous recombination, and spotted onto solid media containing high (2%) and low (0.2 and 0.14%) concentrations of glucose with and without thiamine (**Fig. 2C** and **Supplementary Fig. 1B**). In the absence of thiamine, cells expressing wild-type and mutant *aly3* genes formed colonies on high-glucose medium (**Fig. 2C**). However, cells expressing *aly3*(4^th^ A) exhibited a severe proliferation defect similar to *aly3*(ST18A)-expressing cells in low glucose. In contrast, *aly3*(1^st^ A), *aly3*(2^nd^ A) or *aly3*(3^rd^ A) did not suppress proliferation compared to *aly3*^+^. These results indicated that among the 18 hydroxylated amino acids in Aly3, 4 Ser/Thr residues clustered near the C-terminus (S582, S584, S585, and T586) were required for cell proliferation in low glucose. As these residues are reportedly phosphorylated and *aly3*(4^th^ A) mutation led to a growth defect in low glucose, similar to TORC2-defective mutations, these residues are likely phosphorylated via the TORC2 pathway.

Next, we asked whether all four of these residues had to be simultaneously phosphorylated for cell proliferation in low glucose or whether phosphorylation at one of them is sufficient. To address this question, we prepared a set of mutant *aly3* alleles, in which A582, A584, A585 or A586 in the *aly3*(4^th^ A) gene was reverted back to the original serine or threonine residue (*aly3*(4^th^ A;A582S), *aly3*(4^th^ A;A584S), *aly3*(4^th^ A;A585S), and *aly3*(4^th^ A;A586T), respectively) (**Fig. 3A**). Additionally, *aly3*(S582A), in which only S582 was changed to alanine, was also prepared. These mutant *aly3* genes were placed under the thiamine-repressible *nmt1* promoter and integrated into the genome of *aly3Δ* cells. Cells harboring the mutant *aly3* were spotted onto high- and low-glucose solid medium with and without thiamine. In the presence of thiamine, all strains proliferated comparably in both high and low glucose (**Supplementary Fig. 1C**). In the absence of thiamine, cells expressing the wild-type and the mutant *aly3* genes proliferated comparably in high glucose (**Fig. 3A**). In low glucose, interestingly, cells expressing *aly3*(4^th^ A;A582S), *aly3*(4^th^ A;A584S), and *aly3*(4^th^ A;A585S) genes proliferated as did *aly3*^+^-expressing control cells, whereas cells expressing *aly3*(4^th^ A;A586T) exhibited a proliferation defect similar to those expressing *aly3*(4^th^ A) or *aly3*(ST18A). Expression of *aly3*(S582A) also enabled cell proliferation in low glucose. These results indicate that S582, S584, and S585 of Aly3 function redundantly. Phosphorylation of Aly3 at any one of these serine residues may be sufficient for cell proliferation in low glucose.

**Figure 3.**
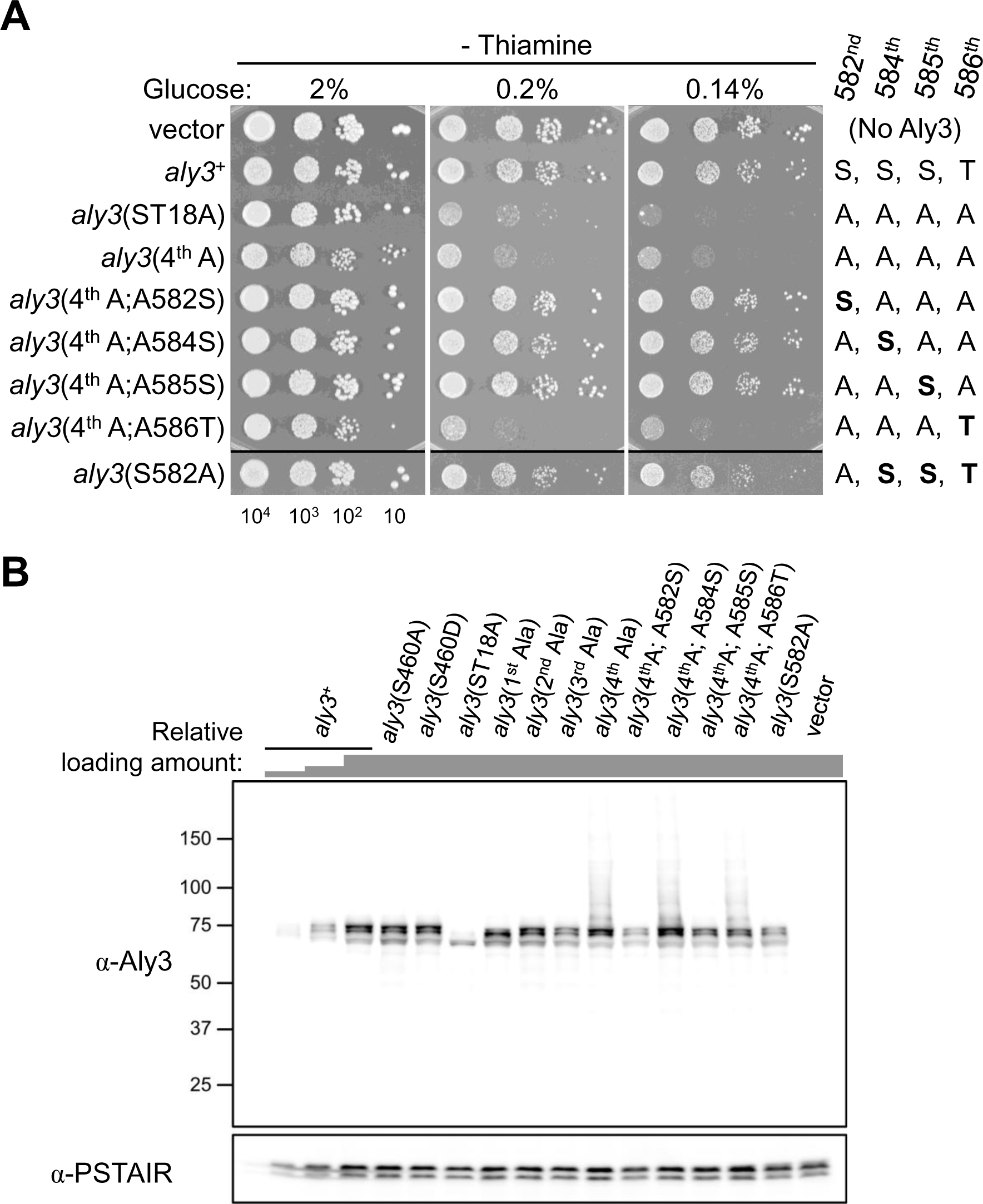
Phosphorylation of Aly3 at one of three serine residues at the C-terminus is required for cell proliferation in low glucose. **A.** Aliquots of *aly3*Δ cells expressing the vector or the wild-type and the indicated phosphorylation-defective *aly3* genes were diluted 10-fold, spotted onto EMM2 solid media containing high (2%) or low (0.2 and 0.14%) concentrations of glucose, but not thiamine and cultivated for 5 days at 33°C. On the right, amino acid sequences of the C-termini of wild-type and mutant Aly3 proteins are shown. At the bottom, numbers of cells spotted are shown. See **Supplementary Figure 1C** for proliferation of these cells in the presence of thiamine. **B.** Expression levels of Aly3 in cells used in the proliferation assay. Cells were cultivated in EMM2 (2% glucose) without thiamine to induce the *aly3* gene. PSTAIR was used as a loading control. The same amount of total protein (OD_595_ = 0.5) was loaded in each lane, while the wild-type, Aly3-expressing sample was loaded in the leftmost 3 lanes to make a 2-fold dilution series. Note that extensive degradation of Aly3 was not observed, whereas smeared signals were observed for Aly3(4^th^ A), Aly3(4^th^ A;A584S) and Aly3(4^th^ A;A586T).

Phosphorylation may affect stability of Aly3, and the difference in Aly3 abundance may impact cell proliferation on low-glucose media, as shown above. As the *aly3* gene transcribed from the *nmt1* promoter was used in the above-mentioned experiments, its expression level was first compared with that from the authentic promoter. A fission yeast wild-type strain expressing Aly3 from the authentic locus (WT *aly3*^+^) and the strain in which the *nmt1* promoter-driven *aly3*^+^ gene was inserted into the genome of *aly3Δ* cells (*P_nmt1_-aly3*^+^) were cultivated in the absence of thiamine to extract total proteins, which were examined by SDS-PAGE. To detect Aly3, we generated a polyclonal anti-Aly3 antibody, as epitope tagging at the either end of Aly3 greatly impaired its function (**Supplementary Fig. 2A-D**). While Aly3 protein in *P_nmt1_-aly3*^+^ was detected as multiple bands in all parts of a 2-fold dilution series, Aly3 in WT *aly3*^+^ was below detectable levels (**Supplementary Fig. 3**), indicating that Aly3 expression levels are much higher in the strains used in this study than in the WT strain, in which the *aly3*^+^ gene was nevertheless transcribed and its gene product was detected in proteomics studies (Marguerat et al., 2012; Carpy et al., 2014; Toyoda et al., 2021).

We next measured expression levels of Aly3 in strains used in the proliferation assay. For this purpose, lysates of cells in which the *aly3* gene was induced, were resolved by SDS-PAGE, and Aly3 was detected by immunoblotting (**Fig. 3B**). Wild-type Aly3 protein was detected as multiple bands ranging from ∼65-80 kDa, which were specific, as no such bands were observed in the vector control. Aly3(ST18A) appeared as a single major band at ∼65 kDa, matching the predicted molecular weight of Aly3. This observation indicated that slower migrating bands were phosphorylated species of Aly3. Importantly, all these strains expressed Aly3 at comparable levels, judging from a 2-fold dilution series of the WT Aly3 protein sample (**Fig. 3B**), suggesting that phosphorylation likely does not affect abundance of Aly3. We concluded that availability of phosphorylation sites of Aly3 protein, but not its expression level, accounts for successful or defective cell proliferation of the strains tested in low glucose.

### Expression of phosphorylation-defective Aly3 causes vacuolar localization of Ght5

We then asked how phosphorylation failure of Aly3 affects intracellular localization of Ght5. Cells used in spot-test analyses, which expressed Ght5-GFP at the native locus, were cultivated to log phase in EMM2 medium with thiamine, and expression of *aly3* was induced by transferring these cells to EMM2 medium without thiamine. After *aly3* expression was induced, cells were transferred to low-glucose (0.08%) EMM2 medium, and cultivated for an additional 4 h before imaging of Ght5-GFP by fluorescence microscopy (**Fig. 4** and **Supplementary Fig. 4**). In cells expressing the wild-type *aly3*^+^ gene, Ght5 was observed primarily at the cell surface and also as puncta in the cytoplasm (**Fig. 4**). Cell-surface localization of Ght5, which is required for cell proliferation in low glucose in *aly3*^+^-expressing cells (Saitoh et al., 2015), was consistent with successful cell proliferation in low glucose (**Figs. 2C** and **3A**). Punctate signals of Ght5, which were similar to those observed upon activation of Aly3-dependent vacuolar transport, e.g., nitrogen starvation (Toyoda et al., 2021), might have occurred due to ectopic expression of the *aly3*^+^ gene from the *nmt1* promoter (**Supplementary Fig. 3**). In contrast to *aly3*^+^-expressing cells, cells expressing the *aly3*(ST18A) or *aly3*(4^th^ A) gene exhibited little or no Ght5 signal on the cell surface. Failure of these cells to retain Ght5 on the cell surface was consistent with their defective cell proliferation in low glucose (**Fig. 2C**). Intriguingly, cell-surface localization of Ght5 was observed in cells expressing *aly3*(4^th^ A;A584S) (**Fig. 4**). Similar restoration of cell-surface localization was observed in cells expressing *aly3*(4^th^ A;A582S) and *aly3*(4^th^ A;A585S), but not *aly3*(4^th^ A;A586T) (**Supplementary Fig. 4**). It is noteworthy that all strains exhibiting cell-surface localization of Ght5 proliferated in low glucose. Background-like cytoplasmic GFP signals, which were prominent in cells showing cell-surface Ght5 localization, but not in *aly3*(ST18A)- and *aly3*(4^th^ A)-expressing cells, might have occurred due to out-of-focus signals of cell membrane-localized Ght5-GFP and/or active intracellular transport of Ght5-GFP. As for other strains used in the spot-test analyses, expression of the *aly3*(1^st^ A), *aly3*(2^nd^ A), *aly3*(3^rd^ A), *aly3*(S460A), *aly3*(S460D), or *aly3*(S582A) genes led to cell-surface localization of Ght5 in low glucose. These results indicate that phosphorylation of Aly3 at the C-terminal 582^nd^, 584^th^, and/or 585^th^ serine residues is required for cell-surface localization of Ght5.

**Figure 4.**
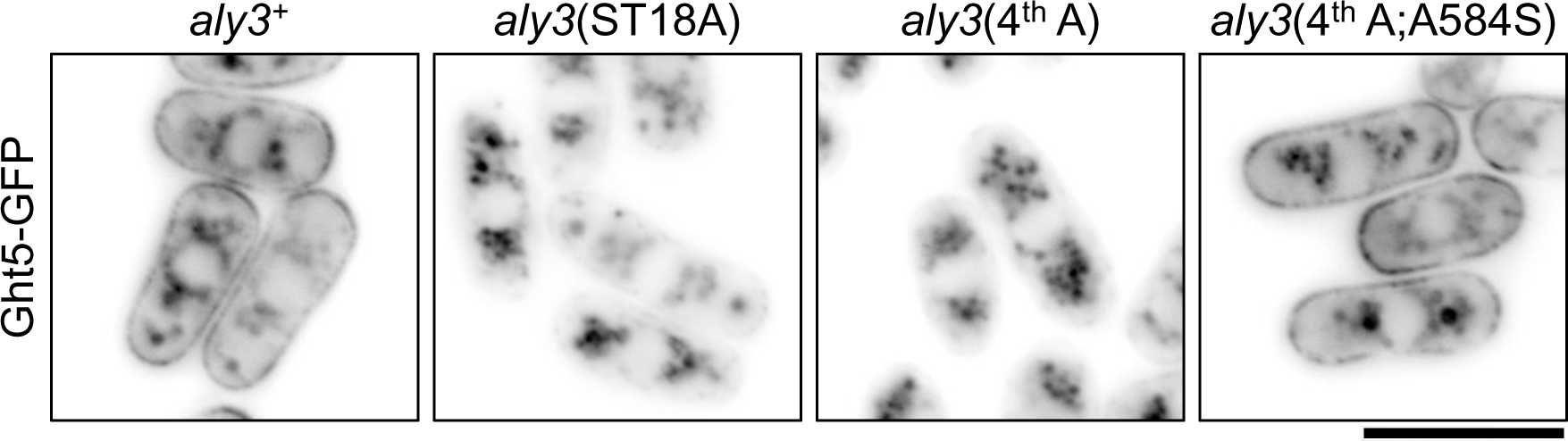
Phosphorylation of Aly3 at the C-terminus is required for cell-surface localization of Ght5. Subcellular localization of Ght5 in living *aly3*Δ cells expressing the wild-type and the indicated phosphorylation-defective *aly3* genes. Cells expressing Ght5-GFP were cultivated in EMM2 (2% glucose) medium without thiamine, and transferred into low-glucose (0.08%) EMM2 medium, and cultivated for an additional 4 h at 33°C. See **Supplementary Figure 4** for Ght5-GFP localization in all strains used in the proliferation assay. Scale bar, 10 μm.

### Phosphorylation of Aly3 at the C-terminus attenuates ubiquitination of Ght5

We then asked how phosphorylation of Aly3 at these C-terminal serine residues ensured cell-surface localization of Ght5. As ubiquitination of Ght5, which was increased in TORC2-defective mutant cells and caused retraction of Ght5 from the cell surface to vacuoles, is dependent on Aly3 (Toyoda et al., 2021), we hypothesized that this phosphorylation inhibited ubiquitination of Ght5. To examine this hypothesis, ubiquitination of Ght5 was assessed. Cells were cultivated in the absence of thiamine to induce the *aly3* gene, and further cultivated in low-glucose medium. Ght5 was affinity-purified from the lysate, and total and ubiquitinated species of Ght5 were detected by immunoblotting (**Fig. 5A**). Affinity-purified Ght5 protein was observed as multiple bands of ∼75 kDa (**Fig. 5A**). As reported previously (Toyoda et al., 2021), the anti-ubiquitin (Ub) antibody detected bands at around 110 and 160 kDa that were not found in the beads-only sample (top panel in **Fig. 5A**). The 110-kDa band, the ubiquitinated species of Ght5, specifically increased in TORC2-defective mutant cells (Toyoda et al., 2021). The extent of ubiquitination of Ght5 was compared among samples by determining relative ratios of the signal intensity of the 110-kDa band to that of the total purified Ght5 (**Fig. 5B**). Expression of Aly3(ST18A) and Aly3(4^th^ A) led to a 3-4-fold increase in Ght5 ubiquitination compared to that of wild-type Aly3 protein. Notably, Aly3(4^th^ A;A584S) restored the level of ubiquitinated Ght5 back to that of wild-type Aly3 protein. Similarly, expression of *aly3*(4^th^ A;A584D), which encodes Aly3(4^th^ A) protein with a phosphorylation-mimetic mutation (Asp) at the 584^th^ residue, also restored the ubiquitination level. These results indicate that phosphorylation of the C-terminus of Aly3 prevents Ght5 from being ubiquitinated, and consequently, being retracted from the cell surface to vacuoles. Phosphorylation at S584 is sufficient to inhibit Ght5 ubiquitination. As cell-surface localization of Ght5 was similarly restored in cells expressing *aly3*(4^th^ A;A582S), *aly3*(4^th^ A;A584S), and *aly3*(4^th^ A;A585S), phosphorylation of Aly3 at S582 and S585 may function redundantly to suppress ubiquitination of Ght5.

**Figure 5.**
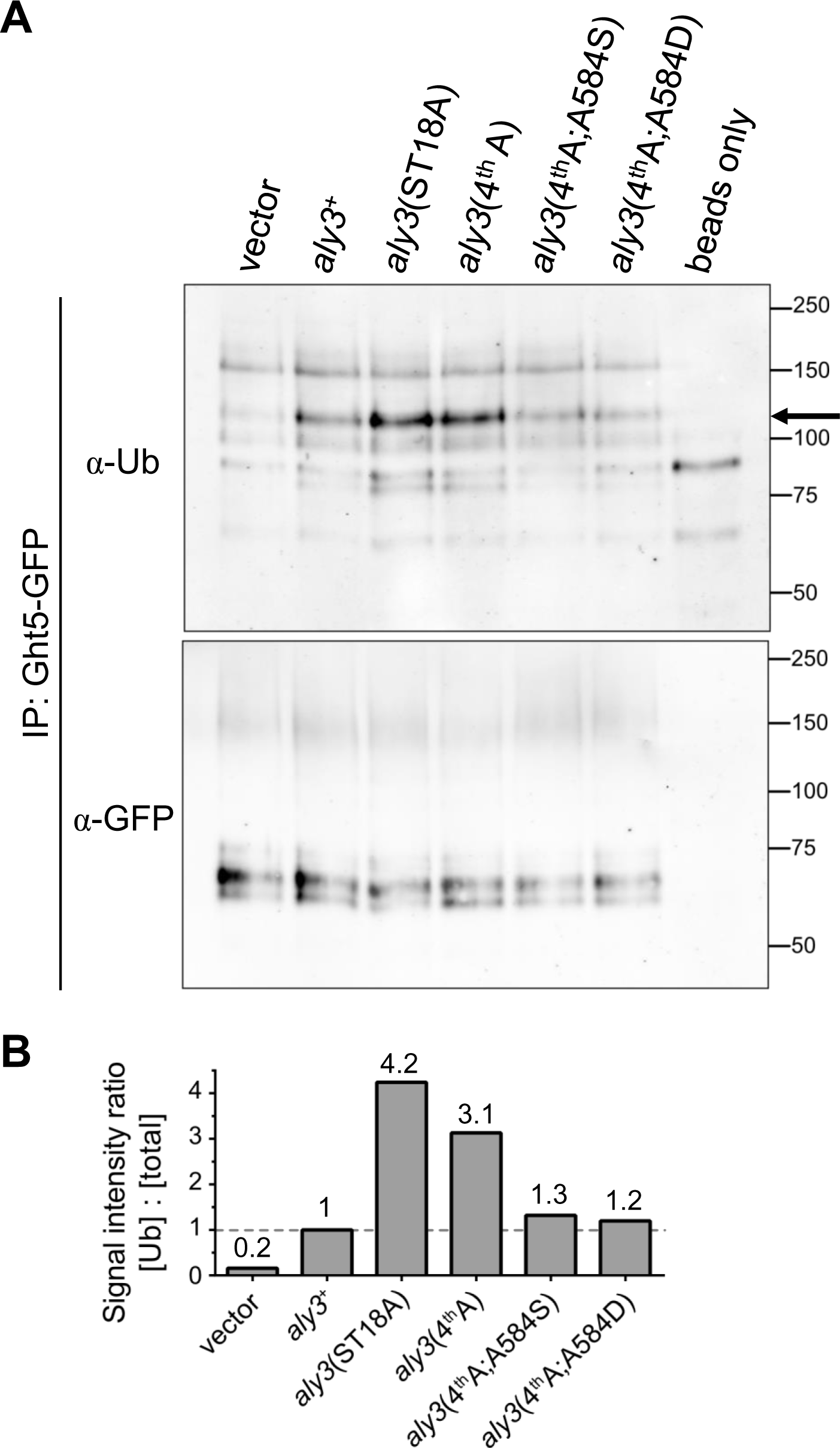
Phosphorylation of Aly3 at the C-terminus reduces ubiquitination of Ght5. **A.** Immunoblot of ubiquitinated and total Ght5-GFP in immunopreciptates. Ght5-GFP in lysates of *aly3*Δ cells expressing the vector or the wild-type and mutant *aly3* genes was affinity-purified using anti-GFP antibody-conjugated beads. Beads only, anti-GFP beads mixed with buffer without lysates. Blots of anti-ubiquitin (α-Ub, top) and anti-GFP (α-GFP, bottom) antibodies are shown as the ubiquitinated species and the total purified Ght5-GFP, respectively. An arrow indicates the 110-kDa band that was more expressed in *aly3*(ST18A) and *aly3*(4^th^ A) cells than in *aly3*^+^-expressing cells. **B.** Quantification of ubiquitinated Ght5-GFP. Signal intensities of ubiquitinated Ght5-GFP ([Ub], the 110-kDa band indicated by the arrow in the α-Ub blot) and the total affinity-purified Ght5-GFP ([total]) were measured, and signal intensity ratios of Ub-Ght5-GFP to total affinity-purified Ght5-GFP are shown in the chart. Signal intensity ratios are normalized using that for *aly3*^+^-expressing cells.

### Aly3 physically interacts with ubiquitin ligases, Pub1 and Pub3

To understand molecular mechanisms by which Aly3 promotes ubiquitination and vacuolar transport of Ght5, we examined the possibility that Aly3 interacts with ubiquitin ligases. In the fission yeast genome, three genes, *pub1*^+^, *pub2*^+^ and *pub3*^+^, encode HECT-type ubiquitin ligases containing the WW motif, which is thought to interact with PY motifs of α-arrestins (Andoh et al., 2002; Tamai and Shimoda, 2002; Harris et al., 2022). Therefore, physical interactions of Aly3 with Pub1-3 proteins were tested by immunoprecipitation. For this experiment, strains expressing *pub1*^+^, *pub2*^+^ and *pub3*^+^ genes that were fused with 13 repeats of a myc epitope at their C-termini and transcribed from their native promoters were produced by homologous recombination. The strain expressing Aly3(4^th^ A) was used as the parental strain, because this strain exhibited more ubiquitination of Ght5 than the wild-type, Aly3-expressing strain (**Fig. 5**). These strains were cultivated in the absence of thiamine to induce the *aly3*(4^th^ A) gene, and further cultivated in low glucose for 4 h. Myc-tagged Pub proteins in cell lysates were affinity-purified. Proteins in the input and affinity-purified (IP) fractions were resolved by SDS-PAGE, and Aly3(4^th^ A) and Pub proteins were detected by immunoblotting (**Fig. 6**). As expected, Pub1-myc, Pub2-myc, and Pub3-myc proteins migrated with their predicted molecular weights (101, 91, and 103 kDa, respectively). In the input fraction, Pub1-myc and Pub3-myc were easily detected, but Pub2-myc appeared as a faint band. Transcription of the *pub2*^+^ gene may be less efficient than that for the *pub1*^+^ and *pub3*^+^ genes, consistent with a previous report (Tamai and Shimoda, 2002). In the IP fraction for the Pub1-myc and Pub3-myc strains, co-precipitation of Aly3(4^th^ A) was clearly detected, indicating physical interactions of Pub1 and Pub3 with Aly3. In contrast, a physical interaction between Pub2 and Aly3 was unclear, because Aly3 signals in the IP fractions for the parental (control) and Pub2-myc strains were similarly faint. Collectively, this result suggests that Aly3 physically interacts at least with Pub1 and Pub3.

**Figure 6.**
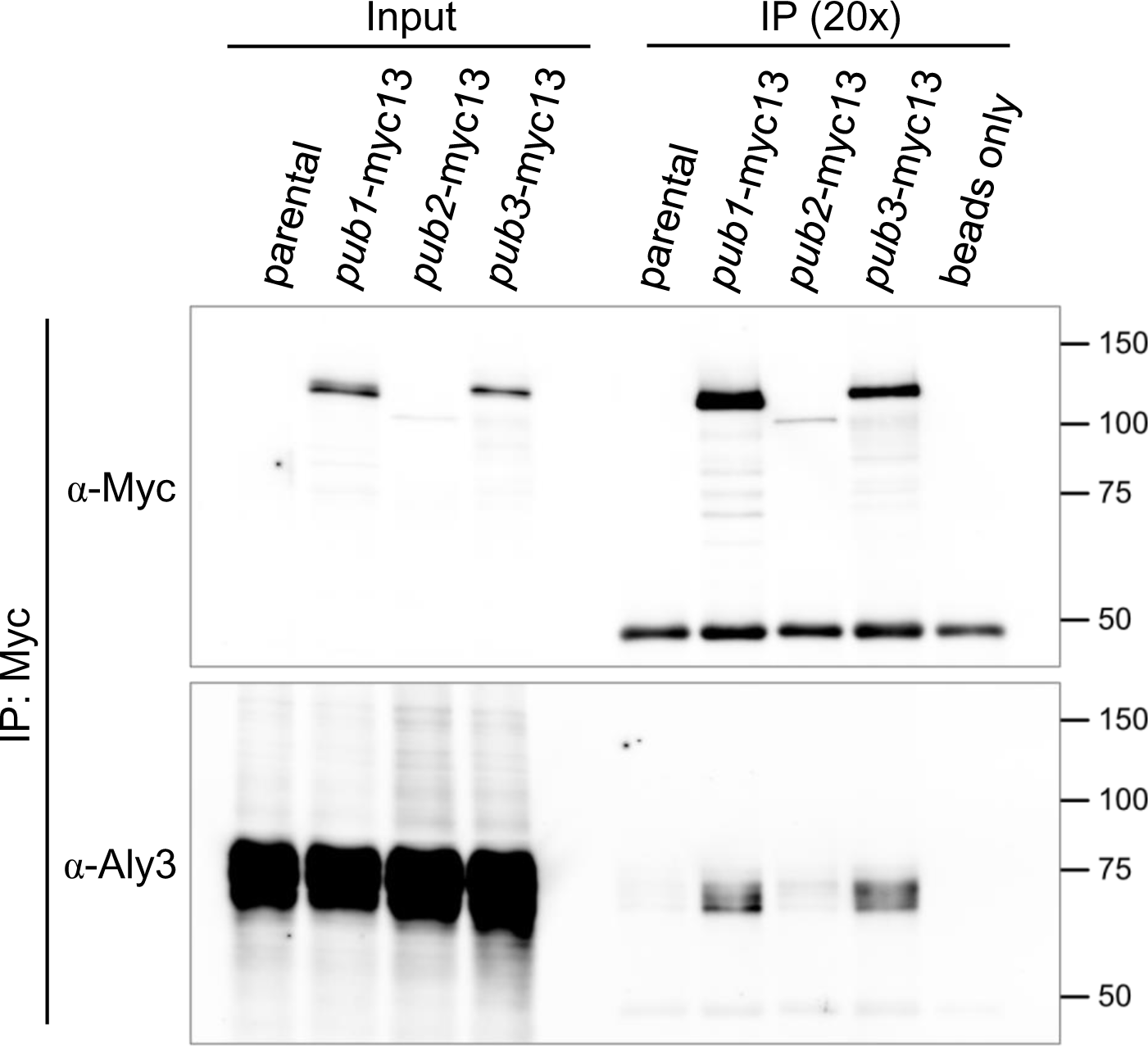
Aly3 physically interacts with ubiquitin ligases Pub1 and Pub3. Immunoblot of Pub1, Pub2, Pub3 and Aly3 in the input and affinity-purified fractions. *aly3*(4^th^ A) strains expressing 13 myc-fused *pub1*^+^, *pub2*^+^ and *pub3*^+^ genes were cultivated in EMM2 medium without thiamine. Myc-tagged Pub proteins were affinity-purified with anti-myc antibody-conjugated beads. The input and immunoprecipitated (IP) fractions were probed with anti-myc (top) and anti-Aly3 antibodies (bottom). The parental strain expressed no myc-tagged proteins. Beads only, anti-myc beads mixed with buffer without lysates. See **Methods** for details.

### *pub3* deletion partly restores proliferation of *gad8^ts^* mutant cells in low glucose

Defects of the *gad8*^ts^ mutant in cell proliferation in low glucose were rescued by deleting the *aly3* gene (Toyoda et al., 2021), because ubiquitination and vacuolar transport of Ght5 was repressed in the *gad8*^ts^ *aly3Δ* mutant. As Aly3 interacts with HECT-type ubiquitin ligases, we examined whether deletion of these ubiquitin ligases, which are likely responsible for ubiquitination of Ght5, could rescue the proliferation defect of the *gad8*^ts^ mutant, similar to *aly3* deletion. *gad8*^ts^ mutant strains lacking the *pub1*^+^, *pub2*^+^ or *pub3*^+^ gene were produced, and spotted on solid YES medium containing different concentrations of glucose (**Fig. 7A** and **Supplementary Fig. 5A-B**). Unexpectedly, the *gad8*^ts^ *pub1*Δ mutant did not proliferate at all, even in high glucose at 33°C, where the growth-defect phenotype on low-glucose media becomes prominent in *gad8*^ts^ mutant cells, and *pub1*Δ single-mutant strains exhibited more severe sensitivity to glucose reduction than the *gad8*^ts^ mutant at 30°C (**Supplementary Figure 5A-B**). Thus, we could not conclusively examine whether *pub1*Δ restores cell proliferation on low-glucose media in *gad8*^ts^ mutant cells. Unlike the *pub1*Δ mutant, *pub2*Δ and *pub3*Δ single-mutants proliferated comparably to the wild type (WT) on both high- and low-glucose solid media at 33°C (**Fig. 7A**). The *gad8*^ts^ *pub2*Δ mutant exhibited proliferation defects in low glucose, as the *gad8*^ts^ mutant did. Unlike *gad8*^ts^ single mutant cells, *gad8*^ts^ *pub3*Δ mutant cells proliferated in low glucose, producing colonies on solid medium containing 0.04 to 0.08% glucose. The restoration of cell proliferation observed in the *gad8*^ts^ *pub3*Δ mutant was somewhat weaker than that in the *gad8*^ts^ *aly3*Δ mutant. The *gad8*^ts^ *pub2*Δ *pub3*Δ triple-mutant cells proliferated in low glucose just like the *gad8*^ts^ *pub3*Δ double mutant, indicating that deletion of the *pub2*^+^ and *pub3*^+^ genes does not synergistically restore cell proliferation of the *gad8*^ts^ mutant in low glucose. Combined with physical interaction data, these results strongly suggest that Aly3 recruits Pub3, but not Pub2, for ubiquitination of Ght5.

**Figure 7.**
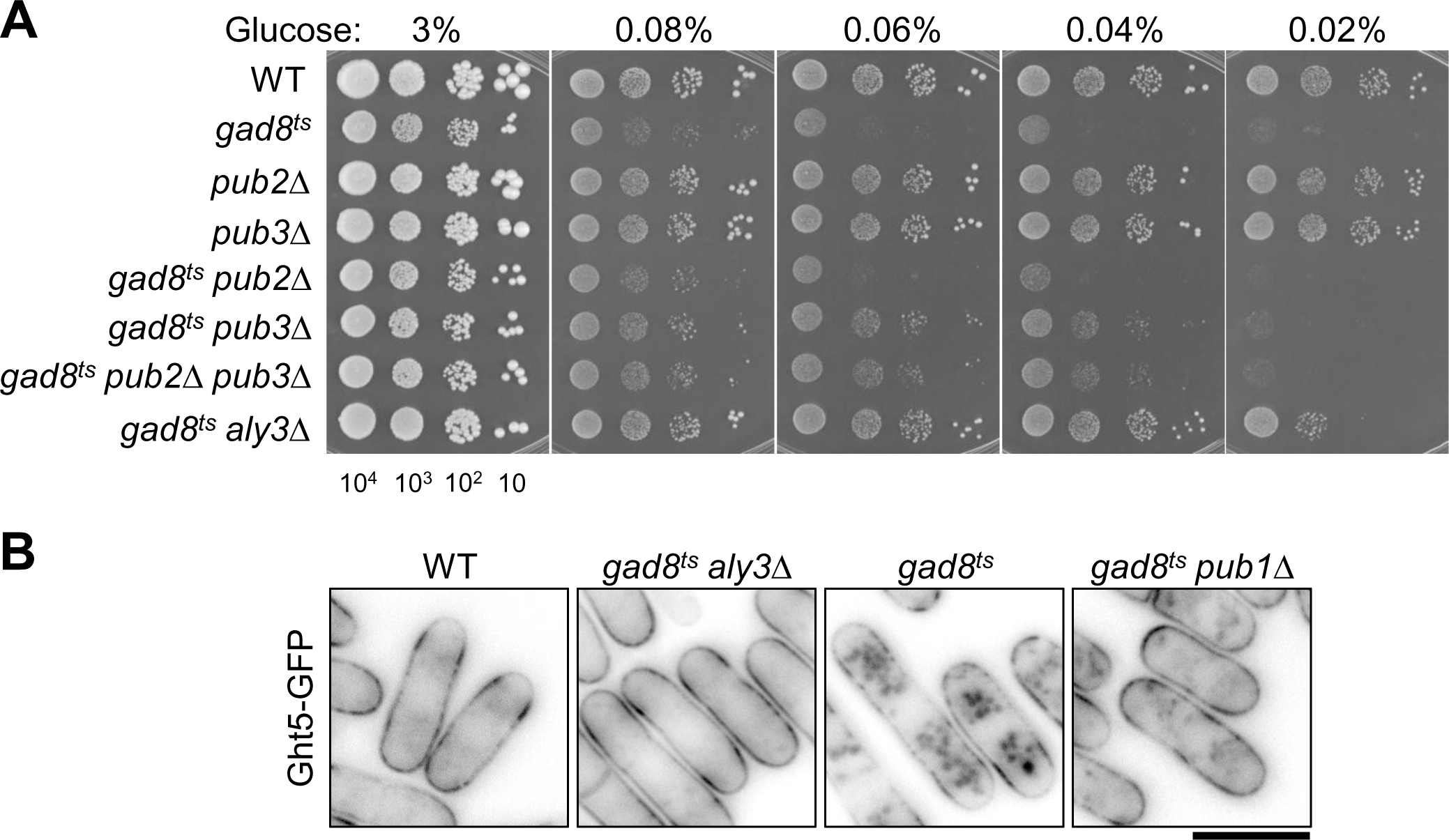
Deletion of Pub1 and Pub3, but not Pub2, suppresses defects of *gad8*^ts^ mutant cells in low glucose. **A.** Aliquots of wild-type (WT) and indicated mutant cells were serially diluted 10-fold, spotted onto YES solid medium containing the indicated concentrations of glucose, and cultivated for 5 days, at 33°C. At the bottom, numbers of cells spotted are shown. See also **Supplementary Figure 5** for proliferation of the *gad8*^ts^ *pub1*Δ mutant. **B.** Subcellular localization of Ght5 in living WT, *gad8*^ts^ mutant, and *gad8*^ts^ mutant cells lacking the indicated genes. Cells expressing Ght5-GFP were cultivated in EMM2 containing 0.08% glucose for 4 h at 33°C. Scale bar, 10 μm.

### *pub1* deletion partly restores cell-surface localization of Ght5 in *gad8^ts^* mutant cells

As genetic interaction between Gad8 and Pub1 could not be examined by colony formation, effects of *pub1* gene deletion on subcellular localization of Ght5 in *gad8*^ts^ mutant cells were assessed. Cells were cultivated to log phase in high-glucose medium at the permissive temperature, 26°C. Then, the medium was replaced with low-glucose medium, and cells were further cultivated at the semi-permissive temperature for the *gad8*^ts^ mutant, 33°C, for 4 h to observe subcellular localization of Ght5. Representative fluorescence images are shown in **Fig. 7B**. As reported previously, Ght5 localized to the entire cell surface in the WT cells, which expressed the *aly3*^+^ gene at the authentic locus of the genome, whereas Ght5 accumulated in the vacuoles in *gad8*^ts^ mutant cells (Saitoh et al., 2015). While vacuolar localization of Ght5 was suppressed in the *gad8*^ts^ *aly3*Δ mutant cells (Toyoda et al., 2021), in *gad8*^ts^ *pub1*Δ mutant cells, vacuolar localization of Ght5 was reduced compared to that in *gad8*^ts^ mutant cells, exhibiting partial restoration. These observations revealed the role of *pub1*^+^ in promoting vacuolar transport of Ght5. Collectively, these results suggest that Pub3 and Pub1, both of which physically and functionally interact with Aly3, promote ubiquitination of Ght5. In addition, the result that deleting either the *aly3*, *pub1*, or *pub3* gene rescued the defects of the *gad8*^ts^ mutant in low glucose is consistent with the idea that Aly3 and its interacting E3 ubiquitin ligases Pub1 and Pub3 together function downstream of the TORC2 pathway under low-glucose conditions.

## Discussion

The present study revealed the molecular mechanism necessary for fission yeast to proliferate in low glucose. Results shown in this and previous studies collectively indicate that phosphorylation of Aly3 at the C-terminus attenuates ubiquitination and subsequent vacuolar transport of the high-affinity glucose transporter, Ght5, ensuring cell proliferation in low glucose (Toyoda et al., 2021). We discuss a model in which Aly3 is regulated via phosphorylation for *S. pombe* cells to adapt to changes in nutrient levels. By comparing our present results to α-arrestin studies of budding yeast and mammals, evolutionarily conserved mechanisms that regulate α-arrestins have become clear.

Based on our present and previous results, we propose that TORC2 pathway-dependent phosphorylation at the C-terminus of Aly3 inhibits ubiquitination of Ght5, ensuring cell-surface localization of Ght5 and cell proliferation in low glucose. Vacuolar transport of Ght5, which is fully repressed by deletion of the *aly3*^+^ gene, is important for chronological life span under low glucose and nitrogen starvation, possibly by acquiring nitrogen via vacuolar degradation of biomolecules (Toyoda and Saitoh, 2021; Toyoda et al., 2021) (**Fig. 8**). Cells expressing the mutant Aly3 protein in which potentially phosphorylatable serine residues are replaced with alanine, i.e., Aly3(ST18A) and Aly3(4^th^ A), exhibited massive accumulation of Ght5 in vacuoles as well as defective cell proliferation in low glucose, suggesting that dephosphorylation activates Aly3 as an adaptor of a ubiquitin ligase for Ght5, which in turn promotes transport of Ght5 from the cell surface to vacuoles. In other words, phosphorylation of C-terminal serine residues of Aly3 (at 582^nd^, 584^th^ and/or 585^th^) inhibits Aly3 for vacuolar transport of Ght5 and ensures Ght5 localization on the cell surface.

**Figure 8.**
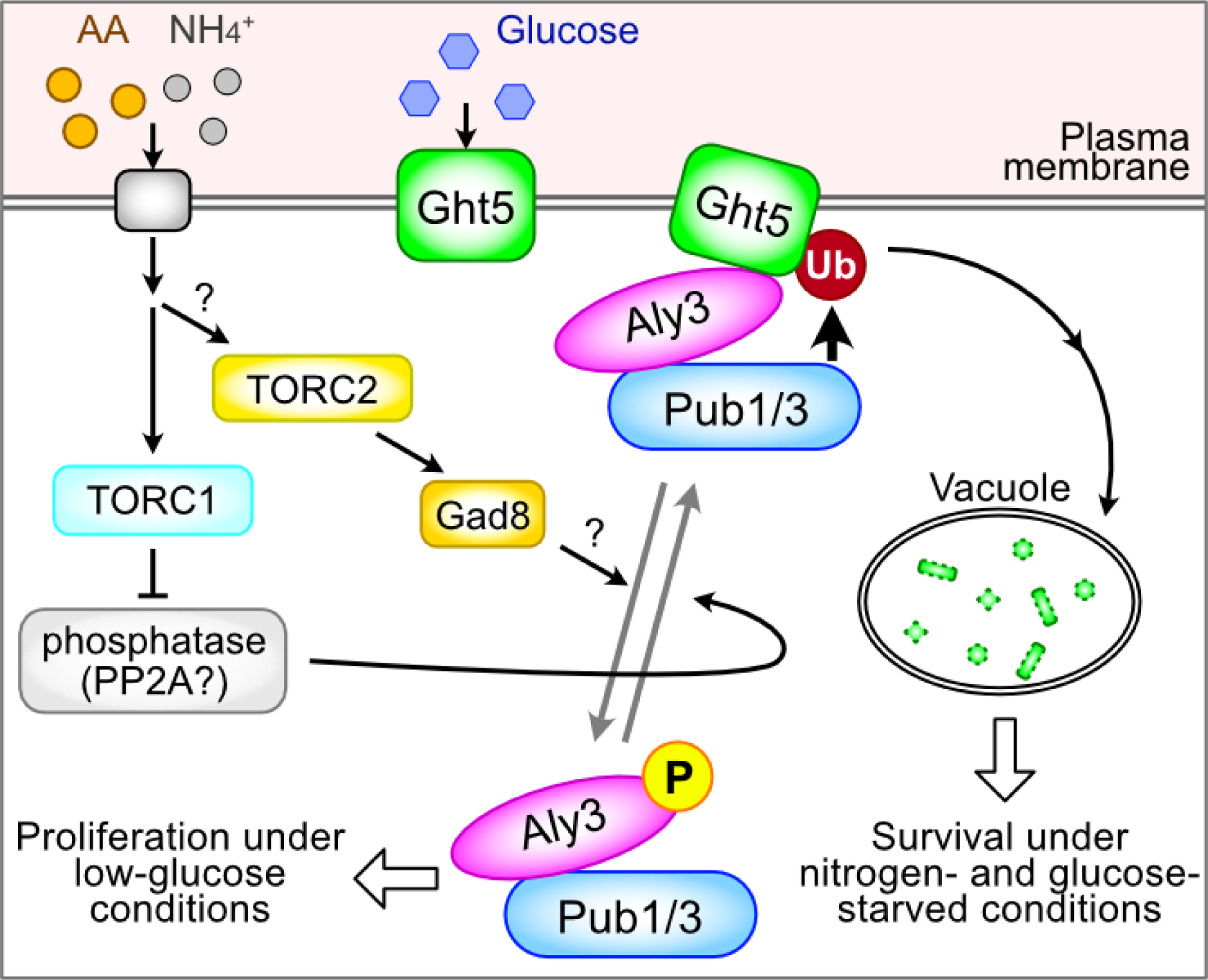
Schematic model showing how Aly3 is controlled by phosphorylation in response to nutritional stress. A model of how Aly3 is regulated by phosphorylation for cellular adaptation to nutritional stress. In the presence of nitrogen, e.g. NH_4_Cl and amino acids, the C-terminal region of Aly3 is phosphorylated in a TORC2 pathway-dependent manner. Upon phosphorylation, Aly3 apparently loses affinity for Ght5, and Ght5 is hypo-ubiquitinated and enabled to localize to the cell surface, which in turn allows *S. pombe* cells to proliferate in low glucose. In contrast, under conditions leading to vacuolar accumulation of Ght5, e.g., nitrogen starvation, the Aly3 C-terminus may be dephosphorylated, possibly due to increased protein phosphatase 2A (PP2A) activity, which is negatively controlled in a TORC1-dependent manner. The net charge of the Aly3 C-terminus then becomes more positive, which may enhance Aly3 interaction with the C-terminal tail of Ght5, which has a negative charge. Pub1 and Pub3 ubiquitin ligases bind to Aly3 and ubiquitinate Ght5 to promote vacuolar transport via the ESCRT machinery. Vacuolar transport of Ght5 maintains cell viability under nitrogen- and glucose-starved conditions. Blue hexagon, glucose; orange and gray circles, amino acids (AA) and ammonium ion (NH_4_^+^), respectively. See main text for details.

Increased ubiquitination and vacuolar localization of Ght5 as well as defective cell proliferation in low glucose observed in TORC2 pathway-deficient mutants were restored by deletion of Aly3 (Toyoda et al., 2021). Essentially the same defective phenotypes caused by expression of Aly3(4^th^ A) were restored by reverting one of the mutated residues back to serine, i.e., Aly3(4^th^ A;A584S). These two analogous results of phenotype restoration suggest that the TORC2 pathway mediates phosphorylation of Aly3 at the C-terminus in low glucose. Interestingly, it was recently reported in a phosphoproteomic study that searched for TORC1 substrates, that phosphorylation of Aly3 at S584 was decreased not only in the presence of Torin1, a potent inhibitor of both TORC1 and TORC2 (Atkin et al., 2014), but also in *tor1*Δ mutant, and that the decreased phosphorylation in *tor1*Δ was largely unchanged, even after Torin1 treatment (Mak et al., 2021). That result suggests that S584 of Aly3 is phosphorylated in a TORC2-dependent manner, consistent with our model. As *S. pombe* TORC2 reportedly activates Gad8 (Ikeda et al., 2008), it is likely that Gad8 phosphorylates Aly3 at S584.

How is phosphorylation of Aly3 controlled according to nutritional status? While Ght5 persists at the cell surface upon glucose limitation in wild-type cells, it is transported to vacuoles upon nitrogen starvation (Toyoda et al., 2021), suggesting that TORC2 is inactivated under nitrogen-starved conditions, and as a consequence, Aly3 is phosphorylated. In a contradictory fashion, the TORC2 pathway is reportedly activated to phosphorylate Gad8 at S546 under nitrogen-starved conditions (Hatano et al., 2015). In contrast, in the presence of nitrogen, TORC1 is active and inhibits the PP2A-like phosphatase, Ppe1, and the absence of nitrogen readily inactivates TORC1 (Hatano et al., 2015; Laor et al., 2015). Therefore, it is more likely that upon nitrogen starvation, the C-terminus of Aly3 is dephosphorylated due to increased phosphatase activity, rather than a decrease in TORC2 kinase activity. Consistent with this idea, deletion of the *tip41*^+^ gene, which encodes a PP2A regulator that enhances phosphatase activity of PP2A (Jacinto et al., 2001; Fenyvuesvolgyi et al., 2005), rescued the surface localization of Ght5 and proliferation of *gad8*^ts^ mutant cells in low glucose (Toyoda et al., 2021). In addition, mutations in negative regulators of TORC1, e.g., Tsc1 and Tsc2, also restored the defects of the *gad8*^ts^ mutant (Toyoda et al., 2021). Activities of PP2A and PP2A-like phosphatases are supposedly controlled via the TORC1 pathways in response to changes in nitrogen concentration, and consequently, the phosphorylation status of Aly3 is determined by the balance between activities of phosphatases and TORC2 kinase.

While phosphorylation of Aly3 at the C-terminus was required for cell proliferation in low glucose, the phosphorylation site(s) was redundant. How does such redundant phosphorylation inhibit Aly3? α-arrestins are thought to interact with a transporter on cell membranes through electrostatic interactions between their own positively charged C-terminal domains and negatively charged cytoplasmic region of the transporter protein (Guiney et al., 2016; Kahlhofer et al., 2021). Indeed, Ght5 has a C-terminal cytoplasmic tail region of around 100 amino acids containing 20 acidic residues plus the 11 phosphorylation sites (Kettenbach et al., 2015). Aly3 has a patch of basic amino acids at the very C-terminus (_580_-RK<u>S</u>L<u>SS</u>TNLVRRGVR*, where an asterisk and an underline indicate the STOP codon and serine residues required for cell proliferation in low glucose, respectively) in addition to those in the arrestin C-terminal domain (_244_-RKEFKRALVKK). These regions may be involved in electrostatic interactions with Ght5. Phosphorylation of serine residues at the C-terminus of Aly3 would neutralize the charge of the C-terminal region, and consequently weaken the interaction between Aly3 and Ght5. Consistently, a non-phosphorylatable mutation in the Aly3 C-terminus, Aly3(4^th^ A), which supposedly prevents neutralization of the charge, caused aberrant ubiquitination and vacuolar localization of Ght5. In addition, phosphorylatable serine residues at the C-terminus (S582, S584, and S585) seem to regulate Aly3 function redundantly, indicating that the net charge of the Aly3 C-terminus, rather than the precise position of the phosphorated serine residue, is important to regulate Aly3 function. Electrostatic interaction between Aly3 and its substrate, Ght5, may be regulated by phosphorylation/dephosphorylation of the Aly3 C-terminus.

Does the phosphorylation status of Aly3 at the C-terminus affect the physical interaction of Aly3 with Pub1/3? It is difficult to answer this question, as fusion with an epitope tag or a fluorescent protein for affinity-purification and visualization compromises the function of Aly3 (**Supplementary Fig. 2**). In budding yeast, Rod1 interacts with Rsp5 regardless of nutritional conditions, indicating that the phosphorylation status of Rod1 does not affect its interaction with Rsp5 (Becuwe et al., 2012). Similarly, Aly3 may interact with Pub1/3 independently of the phosphorylation status of Aly3. As discussed above, it is more likely that C-terminal phosphorylation of Aly3 controls Aly3 interaction with Ght5.

Phosphorylation of α-arrestins constitutes an evolutionarily conserved switch that inhibits endocytosis of hexose transporters (O’Donnell and Schmidt, 2019; Toyoda and Saitoh, 2021). Our studies suggest that the TORC2 pathway, including the Akt-like Gad8 kinase, phosphorylates Aly3 at its C-terminus, whereas *S. cerevisiae* Rod1 and Rog3 are phosphorylated solely by Snf1p/AMPK (Llopis-Torregrosa et al., 2016). In mammals, Akt and AMPK phosphorylate TXNIP at S308 upon insulin stimulation and nutrient stress, including glucose starvation, respectively (Wu et al., 2013; Waldhart et al., 2017), to inactivate TXNIP-dependent endocytosis of GLUT1. Thus, these organisms seem at least partly to share similar mechanisms to control α-arrestins, although the kinases phosphorylating α-arrestins differ between fission and budding yeasts. The mechanisms shared by *S. pombe* and mammals indicate that fission yeast is an excellent model to understand α-arrestin-dependent mechanisms to adapt to nutritional stress in humans. Human α-arrestins, including TXNIP, are associated with progression of diseases such as cancer, diabetes, and Alzheimer’s disease (Zbieralski and Wawrzycka, 2022), although underlying mechanisms are still poorly understood. In the future, identification of the control mechanisms of the 11 *S. pombe* arrestin-related proteins, many of which are still poorly characterized, will contribute to better understanding of α-arrestin-associated human diseases.

## Materials and Methods

### Strains and general techniques

Culture media used for fission yeast cultivation were yeast extract with supplements (YES, rich medium) and EMM2 (minimal medium) (Moreno et al., 1991) with modified concentrations of glucose, as indicated. Unless otherwise stated, wild-type cells were cultivated at 33°C, whereas strains having a mutation in the *gad8*^+^, *pub1*^+^, *pub2*^+^ or *pub3*^+^ gene were cultivated at 26°C. The *S. pombe* strain expressing Ght5-GFP was reported previously (Saitoh et al., 2015). For gene disruption and epitope tagging, a PCR-mediated method was used (Krawchuk and Wahls, 1999). Gene deletion in each strain was confirmed by PCR. To insert the wild-type or mutated *aly3* or *ght5* gene driven by the inducible *nmt1* promoter into the *S. pombe* genome, the pREP1 plasmid expressing the wild-type or mutant *aly3* or *ght5* gene was digested with the restriction enzyme, PacI (Maundrell, 1990, 1993). The digested and purified plasmid was transformed into a leucine-auxotroph strain harboring the *aly3*Δ or *ght5*Δ mutation. Strains that formed colonies on EMM2 solid medium were checked by PCR to isolate strains that had the *nmt1* promoter-driven *aly3* or *ght5* gene inserted in the desired region of the genome, the 3’ UTR region of the *nmt1*^+^ locus. Thiamine (10 μM) was added to EMM2 medium to repress expression of *nmt1* promoter-driven genes, and cells were washed three times in thiamine-free EMM2 medium to induce these genes. To produce a strain expressing the N-terminally tagged 3HA-*aly3* gene driven by the native promoter, a CRISPR/Cas9-based method was used (Hayashi and Tanaka, 2019). A leucine-auxotroph *S. pombe* strain was transformed with the gRNA-Cas9 vector as well as a dsDNA fragment containing a part of 5’ UTR of *aly3*^+^, 3x HA epitope and a part of the *aly3*^+^ coding sequence. Strains that formed colonies on EMM2 solid medium were checked by PCR to isolate strains that had homologous recombination in the desired region. Successfully produced strains were cultivated on YES medium to let them drop the gRNA-Cas9 vector.

### Light microscopy and image analysis

To observe subcellular localization of Ght5-GFP, fission yeast cells were observed with a fluorescence microscope (IX81, Olympus, Tokyo, Japan) equipped with a 100x objective lens (numerical aperture 1.40), and a complementary metal oxide semiconductor (CMOS) camera (ORCA fusion, Hamamatsu Photonics, Japan). To image live cells, the cell culture concentrated by brief centrifugation was kept on ice until use. Brightness and contrast of acquired images were adjusted with ImageJ software (National Institutes of Health, Bethesda, MD, USA).

### Immunoprecipitation, immunoblotting and image quantification

Affinity purification of Ght5-GFP was performed as reported previously (Toyoda et al., 2021). Essentially the same method with slight modifications was used for affinity purification of the 13 myc epitope-tagged Pub1, Pub2 and Pub3 proteins, as detailed below. Cells cultivated in EMM2 medium were washed once in ice cold wash buffer [50 mM Tris-HCl (pH 7.5), 150 mM NaCl, 5 mM NaF, and 1 mM phenylmethylsulfonyl fluoride (PMSF)], and stored at -80 °C until use. Cells were disrupted by vortexing with glass beads in cold lysis buffer [50 mM Tris-HCl (pH 7.5), 150 mM NaCl, 5 mM ethylenediaminetetraacetic acid (EDTA), 10% glycerol, 20 mM β-glycerophosphate, 10 mM *p*-nitrophenyl phosphate, 10 mM NaF, 0.1 mM sodium orthovanadate (Na_3_VO_4_), 0.1 mM dithiothreitol (DTT), 0.5% Nonidet P-40 (NP-40), 10 μM MG-132 (Calbiochem, Darmstadt, Germany), 10 μM ubiquitin isopeptidase inhibitor I G5 (G5, Calbiochem), 1x protease inhibitor cocktail (PIC, Nacalai Tesque, Kyoto, Japan), and 2 mM PMSF]. Cell lysate was centrifuged at 17,700 g for 15 min at 4 °C in a microcentrifuge (Kitman-24, Tomy Seiko, Tokyo, Japan) to prepare input protein extracts for affinity purification. The protein concentration in samples was determined with the Bradford assay using a Protein Assay Dye Reagent (#500-0006, Bio-Rad, Richmond, CA) with optical density reading at 595 nm (OD_595_). The extract with a protein amount of OD_595_=30 was mixed with anti-myc beads (#016-26503, Fujifilm Wako Pure Chemical Corporation, Japan), which had been washed in beads wash buffer [10 mM Tris-HCl (pH 7.5), 150 mM NaCl and 0.5 mM EDTA]. The mixture was incubated on a rotating mixer for 2 h at 4 °C. Anti-myc beads were washed 3 times in immunoprecipitation (IP) wash buffer [50 mM Tris-HCl (pH 7.5), 150 mM NaCl, 5 mM EDTA, 10% glycerol, 2 mM β-glycerophosphate, 1 mM *p*-nitrophenyl phosphate, 1 mM NaF, 0.01 mM Na_3_VO_4_, 0.05% NP-40, 1 μM MG-132, 1 μM G5, 0.1x PIC, and 0.1 mM PMSF]. Anti-myc bead-bound proteins were eluted by boiling for 5 min in sodium dodecyl sulfate (SDS) sample buffer [62.5 mM Tris-HCl (pH 6.8), 5% glycerol, 2% SDS, 2.5% β-mercaptoethanol, and approximately 0.005% bromophenol blue]. One-third the amount of the IP (equivalent to OD_595_=10) and the input (OD_595_=0.5) fractions were resolved by SDS-PAGE, and transferred to a nitrocellulose membrane. To detect HA-tagged and non-tagged Aly3 proteins, crude lysates were prepared as reported previously (Saitoh et al., 2015), using the cold lysis buffer containing 0.2% NP-40 and 1 mM DTT.

Immunoblotting was performed as described previously (Toyoda et al., 2018; Toyoda et al., 2021). Primary antibodies used were anti-GFP (1:2000, rabbit, ab32146, Abcam, Cambridge, UK), anti-HA (1:1000, rabbit, #3724, Cell Signaling Technology, Danvers, MA, USA), anti-ubiquitin (1:200, mouse, MK-11-3, MBL, Nagoya, Japan), anti-Myc (1:500, mouse, sc-40, Santa Cruz Biotechnology, Santa Cruz, CA, USA) and anti-PSTAIR (1:3000, mouse, P7962, Merck, Darmstadt, Germany). Anti-Aly3 polyclonal antibody (1:1000, rabbit) was raised against a part of Aly3 polypeptide (355^th^ to 373^rd^) and affinity-purified. Secondary antibodies used were CF680- or CF770-conjugated anti-mouse and anti-rabbit IgG antibodies (1:5000, #20067 and #20077, Biotium, Hayward, CA, USA). Antibodies were diluted in Blocking One (Nacalai Tesque). Nitrocellulose membranes were washed in Tris-buffered saline (TBS) containing 0.1% Tween-20 (Nacalai Tesque). Fluorescence of membranes was detected on an Odyssey instrument (LI-COR, Lincoln, NE, USA). Fluorescence intensities were analyzed using Image Studio Lite software (LI-COR). To quantify ubiquitinated species of Ght5-GFP (Ub-Ght5-GFP) migrated at 110 kDa in the IP fraction, fluorescence intensity of the 110-kDa band was measured with the surrounding blank area used as a background. Then, the intensity of Ub-Ght5-GFP was divided by that of the total Ght5-GFP in the IP fraction to obtain the signal intensity ratio of Ub-Ght5-GFP to the total Ght5-GFP.

## Acknowledgements

We thank Motohiro Iikura, Yoshino Itoh, and Saho Shirakawa for their help in producing strains. This study was supported by Grants-in-Aid for Scientific Research (C) from the Japan Society for the Promotion of Science (20K06630 and 23K05758 to YT, and 20K06648 to SS).

## Author contributions

Conceptualization: YT, SS; Methodology: YT, SS; Investigation: YT, FM, SS; Writing - original draft: YT, SS; Writing - review & editing: YT, SS; Supervision: SS; Funding acquisition: YT, SS.

---

**Supplementary Figure 1.**
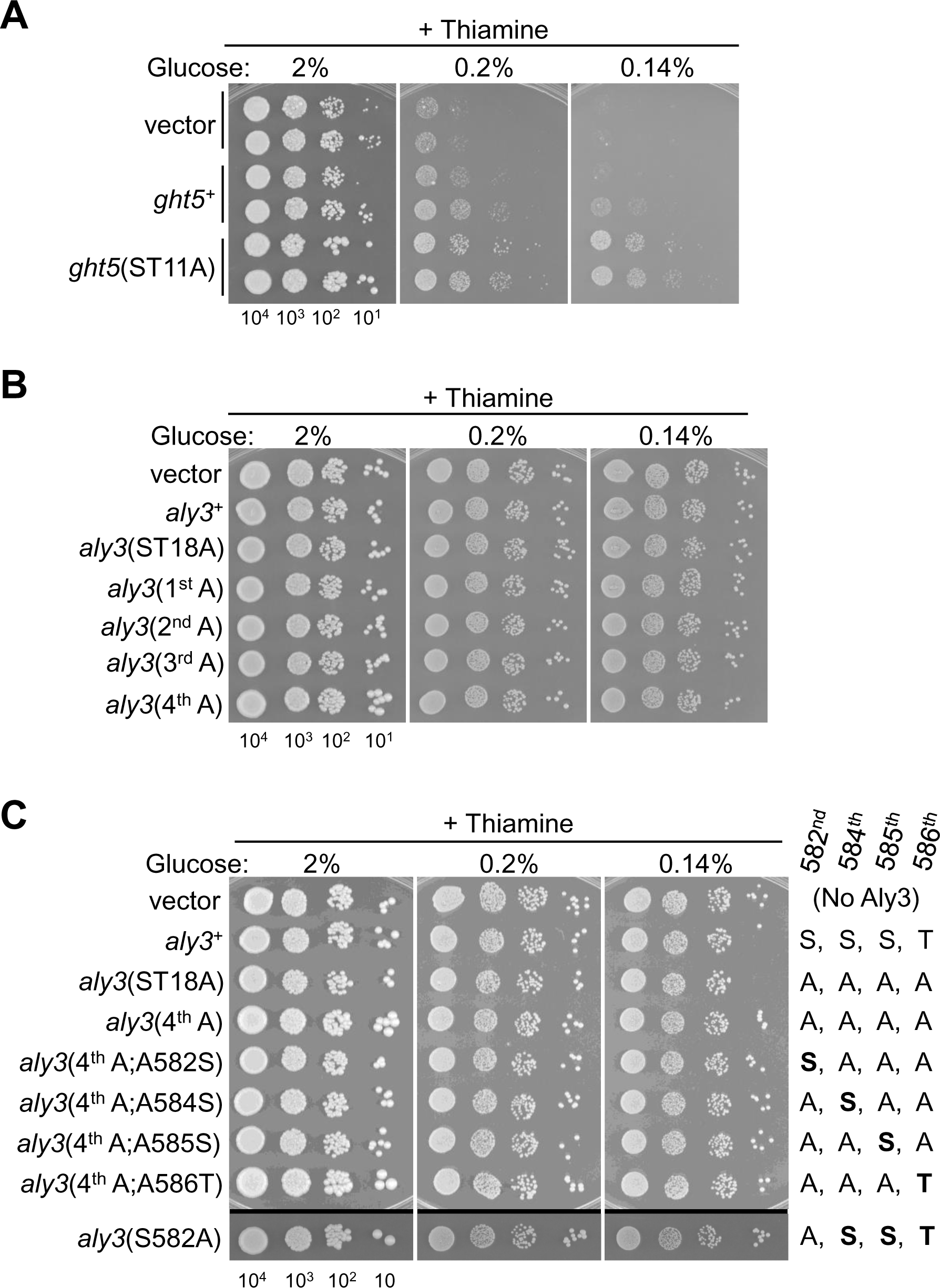
Proliferation of cells expressing phosphorylation-deficient Ght5 or Aly3 protein in the presence of thiamine. **A-C.** Cell proliferation under high- and low-glucose conditions in the presence of thiamine, an expression-repressive condition. Aliquots of cells were serially diluted 10-fold, spotted onto EMM2 solid medium containing thiamine and the indicated concentrations of glucose, and cultivated for 5 days at 33°C. At the bottom, numbers of cells spotted are shown. In panel **A**, *ght5*Δ cells expressing the vector or the indicated gene were used, and two independent clones were tested for each strain. In panels **B** and **C**, *aly3*Δ cells expressing the vector or the indicated gene were used. Note that in panel **A**, under low-glucose (0.2 and 0.14%) conditions, *ght5*Δ cells expressing the *ght5*(ST11A) gene proliferated better than those expressing the *ght5*^+^ gene and the vector. On the right of panel **C**, amino acid sequences of the C-terminus of the wild-type and the phosphorylation-defective Aly3 proteins are shown.

**Supplementary Figure 2.**
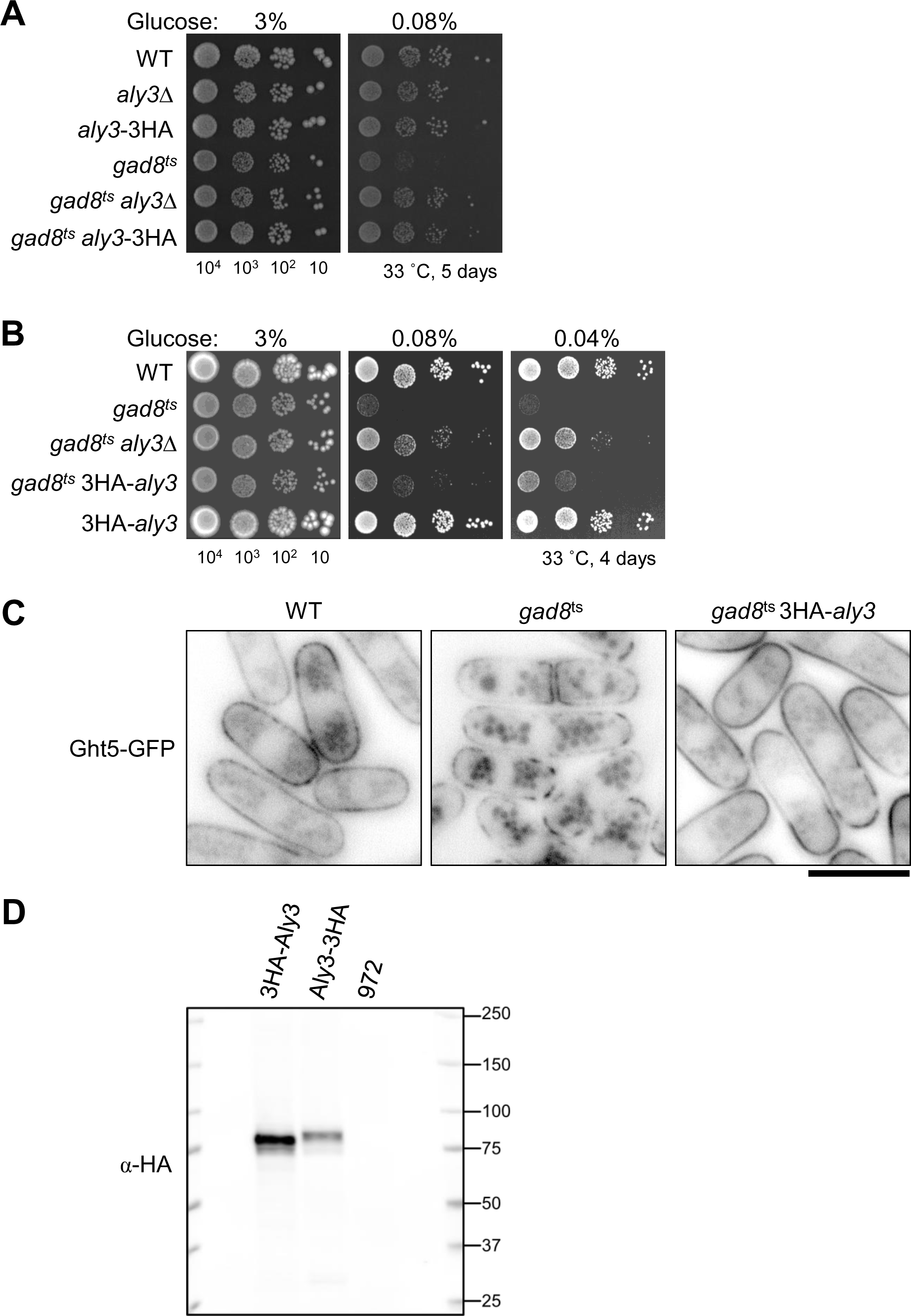
Epitope-tagged Aly3 proteins are not functional. **A-B.** Proliferation of *gad8*^ts^ mutant cells expressing HA epitope-tagged Aly3 proteins from the authentic *aly3*^+^ locus driven by the native *aly3* promoter, in high and low glucose. Aliquots of the wild-type (WT) and indicated strains were serially diluted 10-fold, spotted onto YES solid medium containing the indicated concentrations of glucose, and cultivated for 5 days (panel **A**) or 4 days (panel **B**) at 33°C. At the bottom, numbers of cells spotted are shown. The *aly3*-3HA and 3HA-*aly3* genes encode C-terminally and N-terminally tagged Aly3 proteins, respectively. Similar to *aly3* deletion, epitope tagging at either end of Aly3 rescued the proliferation defects of *gad8*^ts^ mutant cells in low glucose (0.08 and 0.04%). **C.** Subcellular localization of Ght5 in living WT, *gad8*^ts^ mutant and *gad8*^ts^ mutant cells expressing the 3HA-*aly3* gene. Cells expressing Ght5-GFP were cultivated in EMM2 medium (0.08%) for 10 h at 33°C. Note that *gad8*^ts^ mutant cells expressing the 3HA-*aly3* genes exhibited more Ght5 signals on the cell surface than *gad8*^ts^ mutant cells. Scale bar, 10 μm. **D.** Immunoblot of HA-tagged Aly3. Expression of Aly3 protein in these cells used in the proliferation assay (panels **A-B**) was assessed in Western blotting using anti-HA antibody. The same quantity of total proteins (OD_595_ = 0.5) was loaded in each lane. The wild-type strain (972) expressed no tagged proteins.

**Supplementary Figure 3.**
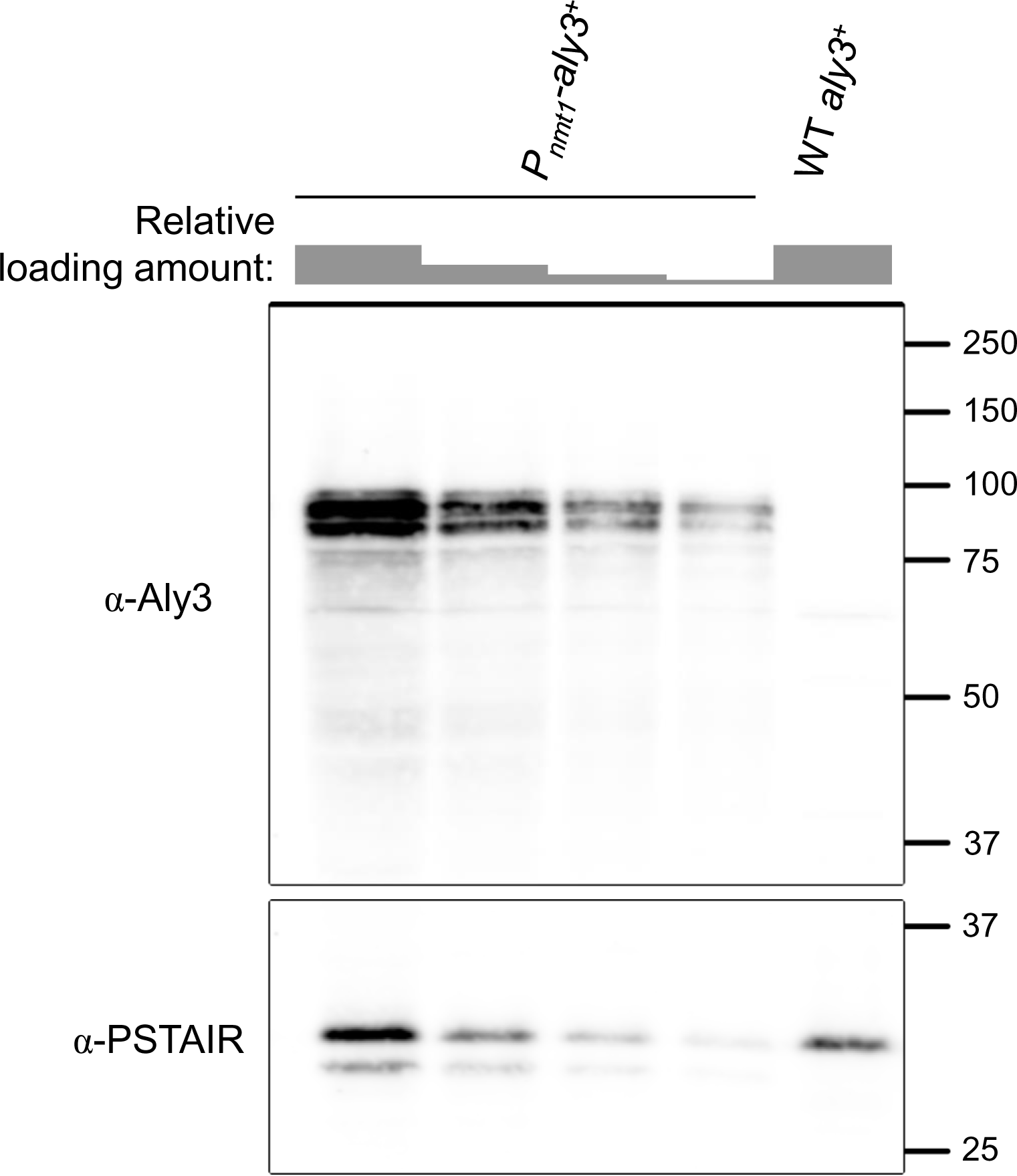
The *nmt1* promoter expresses the *aly3*^+^ gene more efficiently than the authentic *aly3* promoter. Immunoblot of Aly3. *aly3*Δ cells expressing wild-type *aly3* genes from the *nmt1* promoter (P*_nmt1_*-*aly3*^+^) and WT cells (WT *aly3*^+^, identical to the WT strain used in **Supplementary Fig. 2A-C**) were cultivated in EMM2 medium lacking thiamine. Protein extracts were resolved by SDS-PAGE and probed with anti-Aly3 antibody. PSTAIR was used as a loading control. For the P*_nmt1_*-*aly3*^+^, a 2-fold dilution series was made by loading total proteins equivalent to OD_595_ = 0.5, 0.25, 0.125 and 0.0625.

**Supplementary Figure 4.**
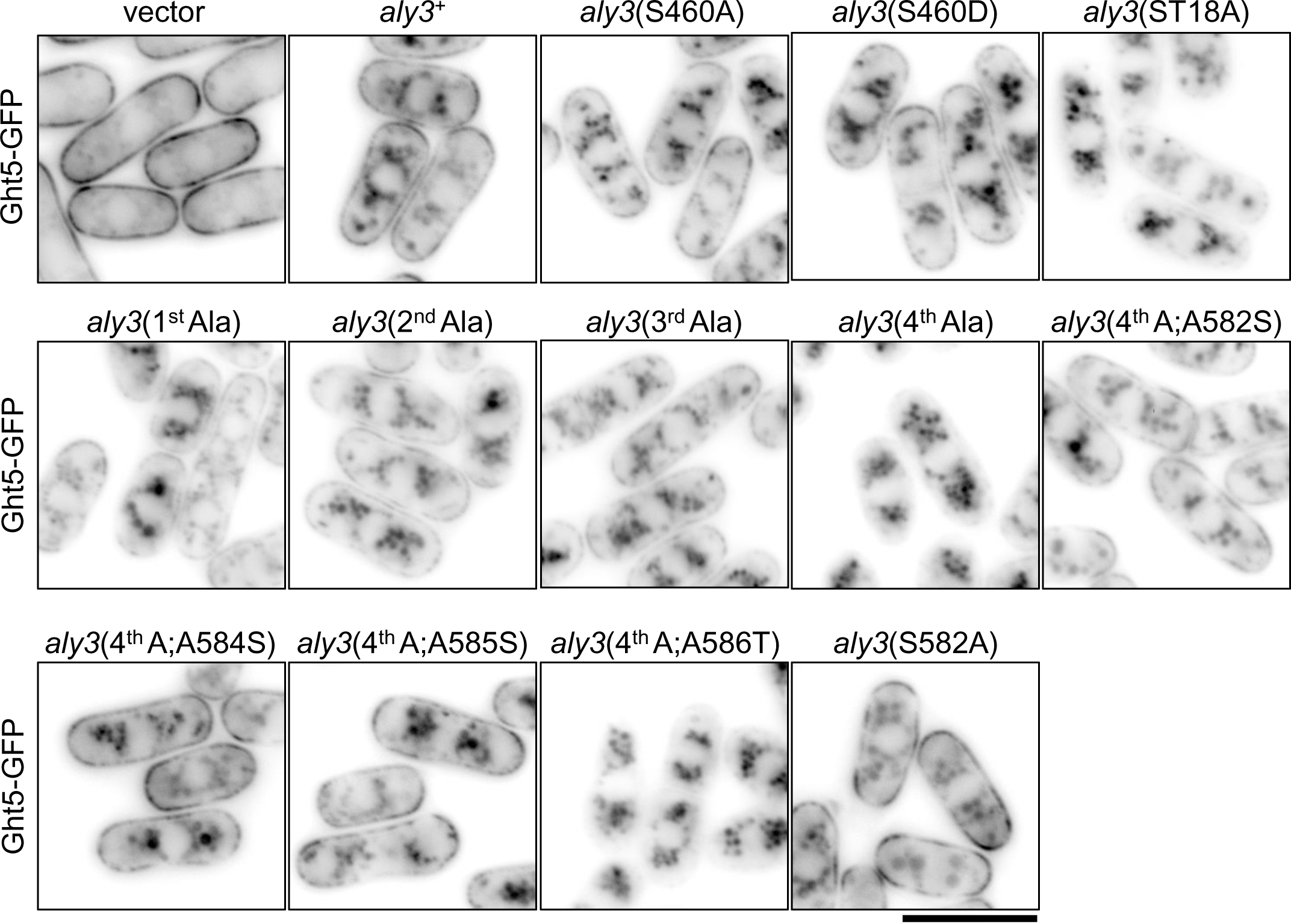
Availability of phosphorylation sites in the C-terminal part of Aly3 accounts for cell-surface localization of Ght5. Subcellular localization of Ght5 in living *aly3*Δ cells expressing the vector or the wild-type and indicated phosphorylation-defective *aly3* genes. Cells expressing Ght5-GFP were cultivated in EMM2 medium (2% glucose) without thiamine to induce *aly3* genes, transferred to low-glucose (0.08%) EMM2 medium, and cultivated for an additional 4 h at 33°C before fluorescence imaging of Ght5. As for Ght5 localization of *aly3*^+^-, *aly3*(ST18A)-, *aly3*(4^th^ A)- and *aly3*(4^th^ A;A584S)-expressing cells, the same micrographs as in Figure 4 are shown. Scale bar, 10 μm.

**Supplementary Figure 5.**
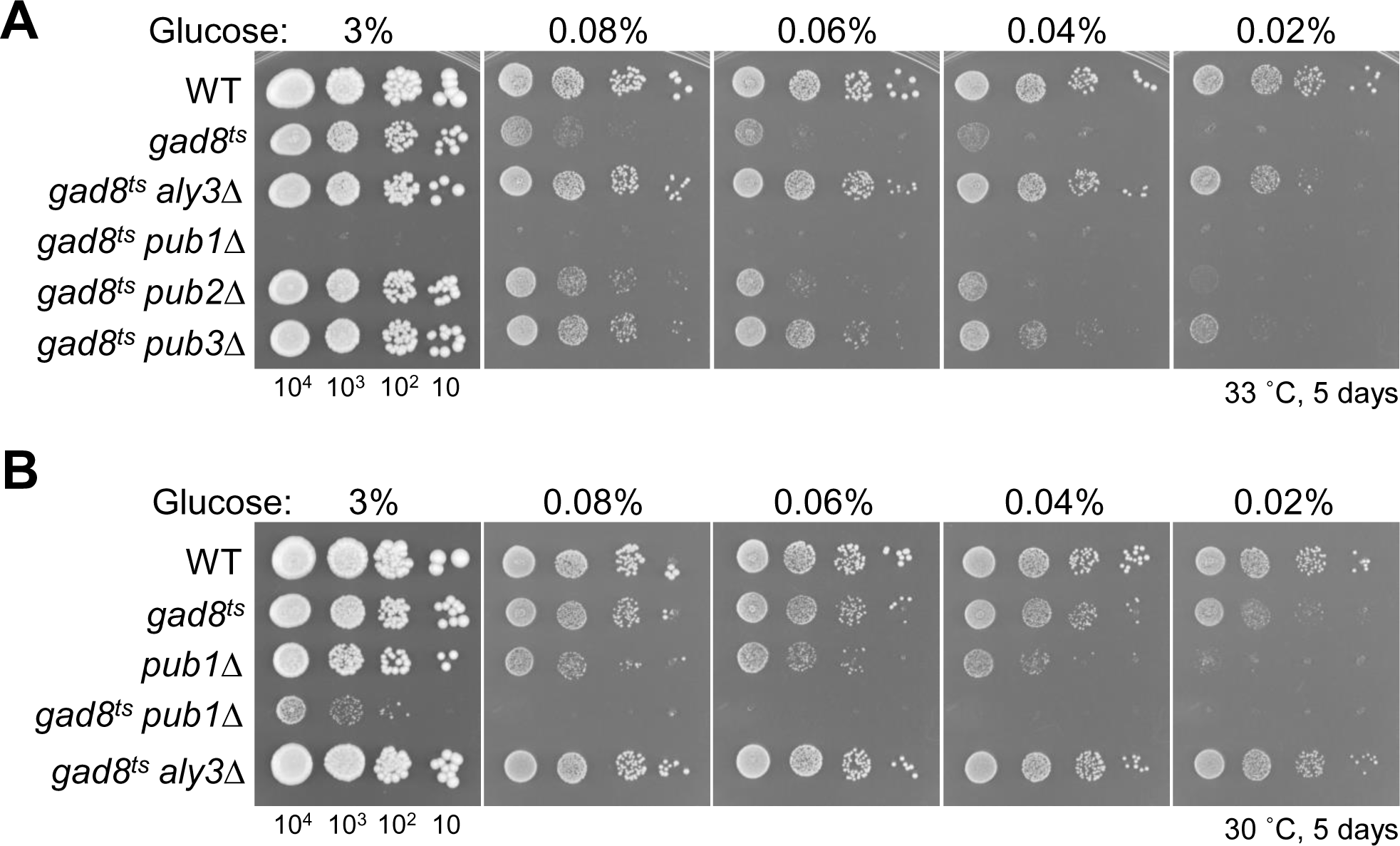
Deletion of Pub3 partially suppresses the proliferation defect of *gad8*^ts^ mutant cells in low glucose. **A-B**. Cell proliferation under high and low glucose. Aliquots of the wild-type (WT) and indicated mutant cells were serially diluted 10-fold, spotted onto YES solid medium containing indicated concentrations of glucose, and cultivated for 5 days at 33°C (panel **A**) or 30°C (panel **B**). At the bottom, numbers of cells spotted are shown. Note that *gad8*^ts^ *pub1*Δ double mutant cells did not proliferate at all at 33°C, and that *pub1*Δ cells exhibited more serious proliferation defects under low-glucose (0.08, 0.06, 0.04 and 0.02%) conditions than *gad8*^ts^ mutant cells.

